# The regulatory potential of transposable elements in maize

**DOI:** 10.1101/2024.07.10.602892

**Authors:** Kerry L. Bubb, Morgan O. Hamm, Thomas W. Tullius, Joseph K. Min, Bryan Ramirez-Corona, Nicholas A. Mueth, Jane Ranchalis, Yizi Mao, Erik J. Bergstrom, Mitchell R. Vollger, Cole Trapnell, Josh T. Cuperus, Andrew B. Stergachis, Christine Queitsch

## Abstract

The genomes of flowering plants consist largely of transposable elements (TEs), some of which modulate gene regulation and function. However, the repetitive nature of TEs and difficulty of mapping individual TEs by short-read-sequencing have hindered our understanding of their regulatory potential. We demonstrate that long-read chromatin fiber sequencing (Fiber-seq) comprehensively identifies accessible chromatin regions (ACRs) and CpG methylation across the maize genome. We uncover stereotypical ACR patterns at young TEs that degenerate with evolutionary age, resulting in TE-enhancers preferentially marked by a novel plant-specific epigenetic feature: simultaneous hyper-CpG methylation and chromatin accessibility. We show that TE ACRs are co-opted as gene promoters and that ACR-containing TEs can facilitate gene amplification. Lastly, we uncover a pervasive epigenetic signature – hypo-5mCpG methylation and diffuse chromatin accessibility – directing TEs to specific loci, including the loci that sparked McClintock’s discovery of TEs.

---

Transposable elements (TEs), first described as ‘controlling elements’ by Barbara McClintock (*1– 7*), have the potential to shape the regulation of the host genome (*8–11*). For example, the insertion of a TE in a regulatory region of the maize domestication gene *teosinte branched1* (*tb1*) enhances its expression, contributing to the increased apical dominance of maize compared to its ancestor teosinte (*12*). Although over 80% of the maize genome is annotated as intact TEs or TE fragments (*13*), a comprehensive analysis of their regulatory potential is lacking. Commonly used methods to map regulatory elements (*i.e.* accessible chromatin regions, ACRs) have relied on short sequence reads which rarely map uniquely within TEs. Here, we use the long-read method Fiber-seq to overcome this limitation and map ACRs across the maize B73 genome. Fiber-seq uses a non-specific DNA *N*^6^-adenine methyltransferase to methylate accessible adenines (*14*) – a modification that is extremely sparse in plants (*15*), including maize (**fig. S1A**) – followed by single-molecule PacBio sequencing of ∼18 kb maize chromatin fibers, enabling the synchronous detection of accessible adenines (m6A) and endogenous cytosine methylation (5mCpG).

## Assessing the single-molecule regulatory landscape of maize with Fiber-seq

We compared Fiber-seq and ATAC-seq using paired samples of leaf protoplasts isolated from 14-day-old dark-grown maize seedlings (**Fig. 1A, fig. S1B-D**). The use of leaf protoplasts minimized cell-type heterogeneity as leaf tissue is enriched in mesophyll cells. We observed that Fiber-seq-derived m6A and 5mCpG calls showed the expected signals at ATAC-seq-derived ACRs and CAGE-defined transcription start sites (TSSs), in addition to the expected correlation of signal intensity with gene expression at TSSs (**fig. S1E-H**). However, unlike ATAC-seq, Fiber-seq also revealed periodic m6A signals downstream of the TSS that were most pronounced for highly expressed genes, reflecting promoter-proximal well-positioned nucleosomes typically measured by MNase-seq (**fig. S1F, G**) (*16*).

**Fig. 1.**
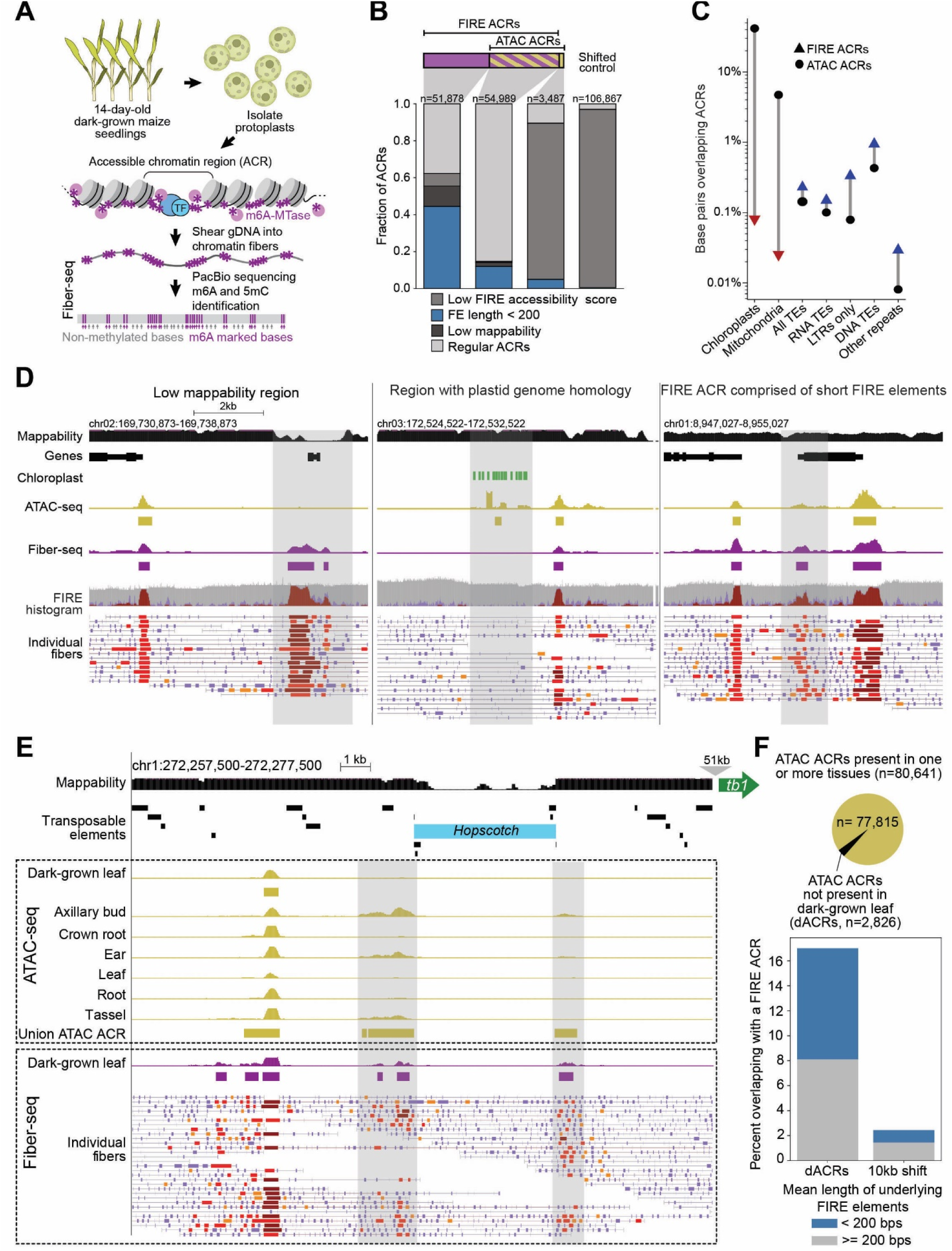
Fiber-seq captures the regulatory landscape of maize comprehensively. **(A)** Experimental scheme. (**B**) ACRs called in paired Fiber-seq and ATAC-seq experiments are shown in three bar graphs representing FIRE ACRs (purple) that did not overlap with ATAC ACRs (n=51,878), FIRE ACRs that overlapped with ATAC ACRs (purple/gold, n=54,989), ATAC ACRs that did not overlap with FIRE ACRs (gold, n=3,487), in addition to a bar graph representing shifted control regions (10kb downstream of FIRE ACRs, n=106,867). ACRs in each category were hierarchically classified as having (i) a FIRE accessibility score of less than 0.25, (ii) a mean FIRE element (FE) length less than 200 bp, (iii) low short-read mappability, or (iv) none of the above (Regular ACR). Stacked bar charts indicate the distribution of these classifiers for each ACR category. (**C**) Percentage of base pairs (Y-axis) overlapping with ACRs called by either Fiber-seq (FIRE ACRs, triangles) or ATAC-seq (ATAC ACRs, circles) for distinct genomic regions (X-axis). As expected, fewer base pairs in regions with homology to plastid and mitochondrial genomic sequence overlap with FIRE ACRs (red triangles) than with ATAC ACRs, while more base pairs in regions annotated as transposable elements or repeats overlap with FIRE ACRs (blue triangles) than with ATAC ACRs. (**D**) Screenshots of three genomic regions illustrating marked differences between FIRE and ATAC ACR calls. Top to bottom, each panel shows the following tracks: genomic location, mappability calculated as in **fig. S2A**, annotated genes, chloroplast sequence, ATAC-seq signal with ATAC ACRs indicated below as rectangles in gold, Fiber-seq signal with FIRE ACRs indicated below as rectangles in purple, FIRE histogram, and individual chromatin fibers with FIRE elements in shades of red, with darker shades indicating greater significance. **Left**, the region highlighted in grey contains two FIRE ACRs but no ATAC ACRs because of low mappability. **Middle**, the chloroplast sequence track indicates high sequence homology at this nuclear locus with the plastid genome. The ATAC ACR in the highlighted region is a false positive due to incorrect mapping of short sequence reads. **Right**, the highlighted region shows a FIRE ACR with underlying short FIRE elements. No ATAC ACR was called. In all panels, ATAC-seq signal is a sliding window histogram displaying the number of Tn5 insertion sites, with the height of each 20 bp bar representing the number of Tn5 insertions within a 150 bp window centered on these 20 bp (minimum=0, maximum=210 mapped reads/16,662,983 total mapped reads=1.26e-5). ATAC ACRs are MACS2 derived peaks (q<0.01) (*49*). Fiber-seq signal is a per-nucleotide average of the scaled, log-transformed ML-model- derived probability of each underlying fiber containing a FIRE element at that nucleotide. Features used by the ML model include m6A density, length of methyltransferase sensitive patch, and A/T content (minimum=0, maximum=100). FIRE ACRs are called by *fiberseq-FIRE* (FDR<0.01) (*50*). FIRE histogram is three overlaid histograms: inferred nucleosome x-coverage (gray), inferred MSP x-coverage (light purple), inferred FIRE element x-coverage (red). Individual fibers are annotated with MSPs (light purple) and FIRE elements (reds, FDR<=5%, and oranges, 5%<FDR<=10%). (**E**) Loci lacking ATAC ACRs in dark-grown maize leaves that show ATAC ACRs in other tissues often overlap with FIRE ACRs comprised of short FIRE elements. A screenshot is shown for the region upstream of the *tb1* gene (green arrow, Zm00001eb054440) in which a hopscotch TE insertion (light blue) generated an enhancer. Top to bottom, tracks are genomic locus, mappability as in **fig.S2A**, in first dotted box, ATAC-seq data: ATAC-seq signal (gold) for dark-grown leaves (this study), subsequent tracks pseudo-bulked single-cell ATAC-seq signal for indicated tissues (*50*), last track, union ATAC ACRs (present in at least one tissue) as golden rectangles, in second dotted box, Fiber-seq data: Fiber-seq signal (purple) in dark-grown leaves and indicated below FIRE ACRs as purple rectangles, individual fibers with short FIRE elements in shades of red. In dark-grown leaves, ACRs were detected by Fiber-seq but not ATAC-seq in the loci flanking the hopscotch TE. However, ATAC ACRs were detected in these loci in axillary bud, ear, and tassel tissue. (**F**) Of the 80,641 union ATAC ACRs across these seven tissues, 2,826 were not detected in dark-grown leaves (differentially accessible ACRs, dACRs, see Methods for details). About 17% of the loci overlapping with these differentially accessible ACRs overlap with FIRE ACRs, and about half of these are FIRE ACRs comprised of short FIRE elements. Statistical analyses and p-values for Fig. 1B and 1F are in **table S1**.

To rigorously distinguish regions with elevated exogenous m6A signal (methyltransferase-sensitive patches, MSP) due to nucleosome linkers from regions representing ACRs (**fig. S1G**), we called FIRE elements (**F**iber-seq **I**nferred **R**egulatory **E**lements) with the semi-supervised machine learning classifier *fiberseq-FIRE* (*17*). After recalibrating *fiberseq-FIRE* for maize, 4.6 million methyltransferase-sensitive patches were classified as actuated FIRE elements (precision >0.9), with the remaining 150 million classified as nucleosome linkers. By aggregating single-molecule FIRE elements across the genome, we called 106,867 FIRE ACRs (FDR <0.01, **Fig. 1B**, **tables S2, S3)**. In contrast, we called only 51,817 ACRs with ATAC-seq (q-value <0.01, t**able S3**), consistent with Fiber-seq revealing a more comprehensive regulatory landscape of maize. Fiber-seq identified the vast majority of ACRs called with ATAC-seq in paired samples (**Fig. 1B**), added ACRs in repeat regions with low mappability, and corrected for false-positive ATAC ACRs, such as those in nuclear genomic regions with homology to plastid or mitochondrial genomes (*18*) (**Fig. 1C, D**, **tables S4, S5**). Signal intensity at ACRs in the paired bulk ATAC-seq strongly correlated with the Fiber-seq signal (**fig. S1I**). However, for a set of ∼40,000 shared ACRs, most were detected with Fiber-seq on half or more of the sequenced chromatin fibers, whereas fewer than 5% of cells showed Tn5 insertions in these ACRs in single-cell ATAC-seq (*19*) (**fig. S1J, table S6**). This comparison illustrates the limitations of single-cell ATAC-seq as a quantitative measure of per-molecule chromatin accessibility. Taken together, our results show that Fiber-seq accurately captures chromatin accessibility and 5mCpG in maize, with single-molecule and single-nucleotide precision, at a sensitivity twice that of ATAC-seq.

## FIRE ACRs comprised of short FIRE elements can mark ATAC ACRs detected in other tissues

The nearly twice as many ACRs identified by Fiber-seq as compared to the paired ATAC-seq ACRs were only in part explained by low mappability of ATAC-seq reads (**fig.S2A, B**). Rather, over half of the FIRE ACRs missed by ATAC-seq were comprised of short FIRE elements (<200 bp) (**Fig.1B, fig. S2C, D**). These FIRE ACRs shared the features typical of ACRs comprised of long FIRE elements identified by both methods such as enrichment of the 6mA signal, depletion of the 5mCpG signal (**fig. S2E**) and genomic distribution (**fig. S2F**).

We detected ACRs comprised of short FIRE elements flanking the TE insertion that introduced a *tb1* enhancer (Studer et al, 2011) but failed to detect these by ATAC-seq (**Fig 1E**). However, these flanking regions were detected as ATAC ACRs in embryonic and reproductive tissues (axillary bud, tassel, and ear) (*19*), suggesting that ACRs comprised of short FIRE elements may mark genomic loci with tissue-specific chromatin accessibility in maize (**Fig. 1E**). To systematically evaluate this possibility, we identified ATAC ACRs present in one or more tissues (*i.e.,* union ACRs) (*19*), and then filtered for the subset of these union ACRs for which the corresponding genomic loci showed only background ATAC signal in dark-grown leaves (*i.e.,* differential ACRs not present in dark-grown leaves, dACRs), yielding 2,826 dACRs from the total of 80,641 union ACRs (**tables S7, S8**). Of the 2,826 dACRs, 480 overlapped with a FIRE ACR (17%), and over half of the 480 overlapped with FIRE ACRs comprised of short FIRE elements (251/480, **Fig. 1F**). This result is thus consistent with these ACRs composed of short FIRE elements corresponding to functional regulatory elements in maize that display tissue-selective activity.

## Distinctive patterns of ACRs mark functional LTR retrotransposons

We next sought to interrogate ACRs in TEs, focusing on long terminal repeat (LTR) retrotransposons because of their prevalence in the maize genome (74.4%) (*19*). Intact LTR retrotransposons are class I TEs with bilateral LTRs that flank an internal region (**Fig. 2A, B**). Each of the bilateral LTRs are thought to contain the regulatory elements, promoters and adjacent enhancers, that drive expression of the TE genes encoded in the internal region (*20*). LTR retrotransposons mobilize through reverse transcription of their mRNA and integration of the cDNA into another genomic location. They are divided into autonomous (which encode the proteins needed for transposition) and non-autonomous (which require proteins encoded by other elements for transposition) LTR retrotransposons. It has been challenging to determine the functional activity of individual LTR retrotransposons because their high sequence identity limits the ability of short-read data to be uniquely mapped to individual LTR retrotransposons (*9*, *11*, *20*).

**Fig. 2.**
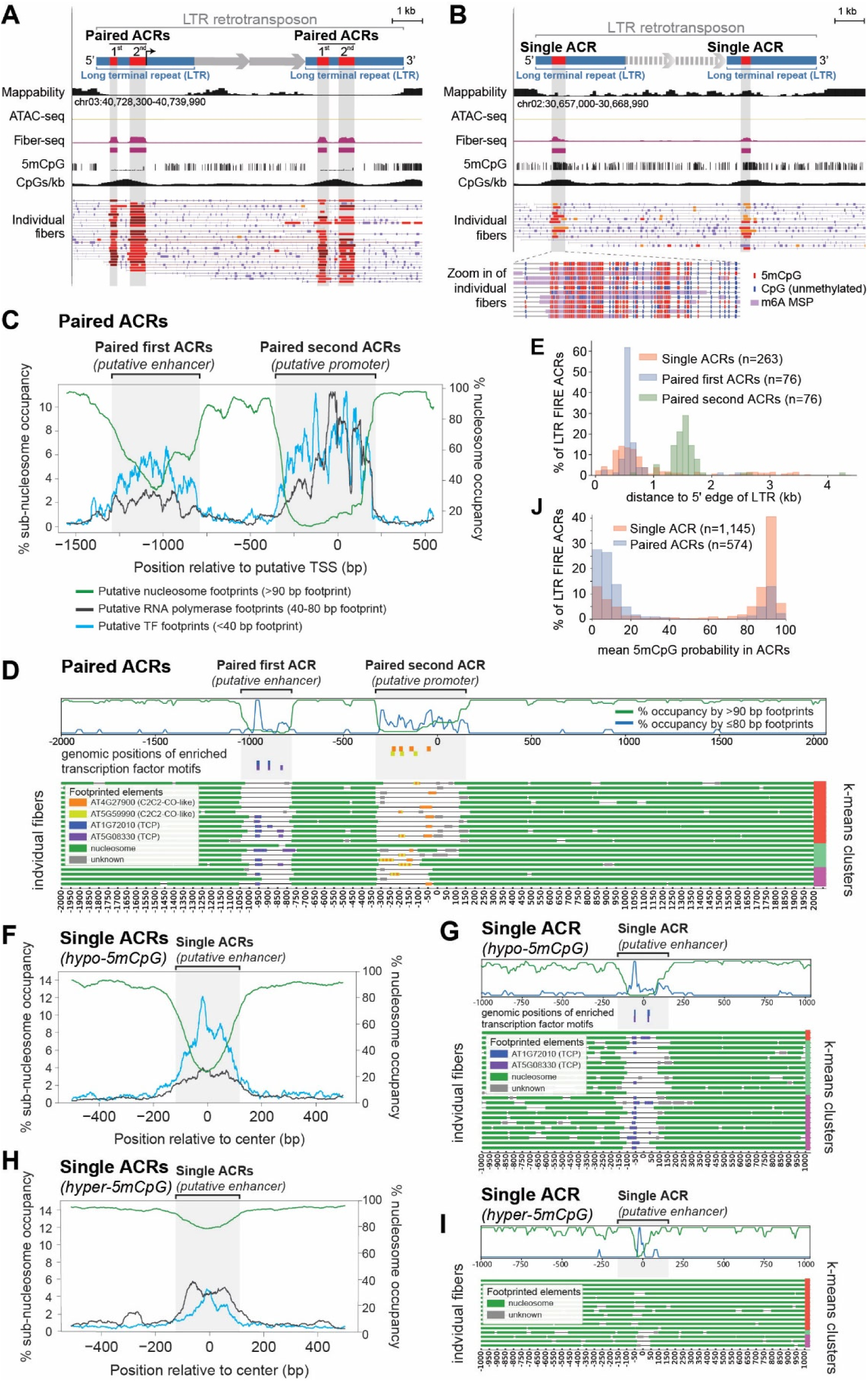
FIRE ACRs in intact LTR retrotransposons identify functional retrotransposons. **(A)** Representative example of an intact LTR retrotransposon with paired FIRE ACRs both in the left and the right long terminal repeats (paired bilateral ACRs in LTRs, ID=LTRRT_14411). Top indicates left and right LTR in blue and paired bilateral FIRE ACRs in red. First and second paired ACRs are labeled, representing the putative enhancer and the putative promoter, respectively. Putative transcription start site is indicated with black arrow, and genes in internal region are indicated. Top to bottom, screenshot shows tracks for genomic location, mappability calculated as in **fig. S2A**, ATAC-seq signal in gold, Fiber-seq signal in purple with FIRE ACRs indicated as purple rectangles below, 5mCpG methylation as the per-CpG methylation probability calculated by pb-CpG-tools (*51*).The vertical axis of the 5mCpG track represents the methylation probability at individual CpG sites, expressed as a percentage, minimum=0, maximum=100. The CpGs/kb track represents a sliding window histogram displaying the number of CpG dinucleotides, with the height of each 100 bp bar representing the number of CpG dinucleotides within a 1 kb window centered on these 100 bp. Paired bilateral ACRs in LTR retrotransposons tended to be hypo- methylated as expected for accessible regions. **(B)** Representative example of an intact LTR retrotransposon with one FIRE ACR both in the left and the right LTR (single bilateral ACRs in LTRs, ID=LTRRT_8308). Tracks as in (**A**). The single FIRE ACRs in this LTR retrotransposon showed high levels of 5mCpG methylation coinciding with the m6A signal, magnified detail below shows methylated 5mCpGs (red) and unmethylated CpGs (blue) and m6A methyltransferse- sensitive patches (purple) on individual fibers. (**C**) Plot depicting the aggregate percent- occupancy of nucleosome-sized footprints (>90bp, green), putative polymerase-sized footprints (40-80bp, dark grey), and putative TF-sized footprints (10-40bp, azure blue) within the putative promoters of all paired ACRs in LTRs. On the X-axis, zero indicates a selected base proximal to the 3’ end of the putative promoter (typically around 75 bp upstream) that is highly conserved across all putative promoters. (**D**) Representative example of footprinted Fiber-seq reads at a paired ACR (zero = chr05:195444369, reverse strand). Top to bottom, (top) the tracks show the count of sub-nucleosomal sized footprints (<80 bp, dark blue) and nucleosome-sized footprints (>90 bp, green) at each position normalized to the maximum count within the region; (middle) blocks showing the genomic position of enriched TF motifs, colored by identity; (bottom) individual Fiber-seq reads with footprints represented by blocks colored by predicted identity (accessible, thin black line; nucleosome, green; unknown, grey; TF, colored corresponding to overlapped motif with multiple overlapped motifs indicated via stripes). Reads are clustered and sorted via k-means clustering with 3 clusters, indicated via a column of colors at right. (**E**) LTR retrotransposons with single bilateral FIRE ACRs tended to maintain the ACRs marking putative enhancers. Histogram of FIRE ACR location relative to the 5’ edge of a given retrotransposon, stratified by type of ACR. (**F**) Plot depicting the aggregate percent occupancy of nucleosome-sized footprints (>90bp, green), putative polymerase-sized footprints (40-80bp, dark grey), and putative TF-sized footprints (10-40bp, azure blue) at positions surrounding the center of all hypo-5mCpG methylated single ACRs. (**G**) Representative example of footprinted Fiber-seq reads at a single hypo-methylated ACR (chr04: 170173642-170173927, forward strand), with position indicated relative to the center of the ACR. Tracks as in (**D**). (**H**) Plot depicting the aggregate percent occupancy of nucleosome-sized footprints (>90bp, green), putative polymerase-sized footprints (40-80bp, dark grey), and putative TF-sized footprints (10-40bp, azure blue) at positions surrounding the center of all hyper-5mCpG methylated single ACRs. (**I**) Representative example of footprinted Fiber-seq reads at a single hyper-methylated ACR (chr04: 206713520-206713627, forward strand), with position indicated relative to the center of the ACR. Tracks as in (**D**), with the motif track omitted due to a lack of motifs. (**J**) Single FIRE ACRs in LTR retrotransposons were more likely to be 5mCpG-methylated than paired bilateral FIRE ACRs, regardless of their position.

Using Fiber-seq, we mapped ACRs residing within each of the 51,882 intact LTR retrotransposons in the maize genome (**table S9**) as well as ACRs in solo LTRs (**fig. S3**, **table S10)**. Only 2% (941/51,882) of intact LTR retrotransposons contained at least one FIRE ACR entirely within one of their bilateral LTRs (**table S9**), consistent with widespread epigenetic silencing of maize LTRs by RNA-mediated DNA methylation, a plant-specific pathway that targets TEs (*21*). Of the 941 ACR-containing LTR retrotransposons, 21% (201/941) contained two adjacent ACRs (paired ACRs, **Fig. 2A, table S9**) in one or both of their LTRs. Furthermore, 94 of these contained the two adjacent ACRs in both LTRs (paired bilateral ACRs, **table S9**).

The paired bilateral ACRs almost always exhibited single-molecule co-accessibility and hypo- 5mCpG methylation (**Fig. 2A**). Given their position relative to the 5’ end of the transposon and the putative transcription start site, these ACRs likely correspond to the putative LTR promoter and enhancer elements (*20*). Consistent with this assumption, the putative promoters showed far higher predicted promoter scores than the putative enhancers when using a validated convolutional neural net model for promoter strength (*23*) (**fig. S4A**). Furthermore, the putative LTR promoter and enhancer elements are enriched in distinct TF motifs, consistent with them having distinct regulatory roles (**tables S11-14)**. To further validate whether the putative LTR enhancers and promoters have distinct regulatory roles, we applied FiberHMM (*23*) to call protein occupancy footprints within them. Prior studies using Fiber-seq show that eukaryotic RNA polymerases display MTase footprints of 40-60bp in size, which is distinct from those of transcription factors (TFs), which are typically <40 bp in size (*11*, *24*). Using this approach, we observed that LTR putative promoters were uniquely marked by MTase footprints of 40-60bp in size (**Fig. 2C**), consistent with these elements being RNA polymerase-bound promoters. In contrast, the LTR putative enhancers were largely marked by MTase footprints of <40 bp in size that were well-aligned with enriched TF motifs (**Fig. 2C, D**), consistent with them being occupied by TFs. The putative enhancer and promoter ACRs were generally separated by 3 or 4 well-positioned mononucleosome footprints, with the regions upstream and downstream of the putative promoter showing larger, less clearly phased di- and trinucleosome footprints, typically associated with heterochromatin (**fig. S5A**). Although the putative enhancer and promoter ACRs were nearly constitutively accessible on all fibers, protein occupancy by 40-60 bp footprints and <40 bp footprints varied substantially between them (**Fig. 2C**).

Together, these findings demonstrate that Fiber-seq enables the identification of individual LTRs within the maize genome that contain active chromatin at both the putative enhancers and promoters, consistent with these LTRs being poised to be functionally active. In total, in maize leaves there are only 94 LTR retrotransposons that contain the functional regulatory elements required for transposon mobilization, with only 76 of these being autonomous LTR retrotransposons.

## LTRs with single ACRs are putative enhancers that display a novel epigenetic signature

Most LTR retrotransposons that harbor an ACR contained only a single ACR in one or both of their LTRs (**Fig. 2B, table S9**). Intact LTR retrotransposons with single ACRs were enriched for containing a single ACR in both of their bilateral LTRs (*i.e.*, single bilateral ACRs, 499/941, 53%). The single ACRs exhibited far greater single-molecule heterogeneity than paired ACRs (**fig. S4B**). Specifically, while paired ACRs showed a bimodal actuation distribution with over half being supported by FIRE elements called in 75% of underlying fibers, only 7% of single ACRs crossed this actuation threshold (**fig. S4B**).

We next examined whether single ACRs preferentially localized to the putative LTR enhancer or to the putative LTR promoter. To accomplish this, we first examined the distance of the ACR to the 5’ LTR edge, which could be measured within the subset of 268 autonomous LTR retrotransposons containing an ACR, as strandedness could be inferred at these sites. We observed that nearly all LTR retrotransposons with a single ACR selectively retained the ACR that positionally corresponds to the putative LTR enhancer element (**Fig. 2E**), suggesting that chromatin accessibility is lost at the position of the putative LTR promoters.

Single ACRs contained predicted TF motifs that were more similar to those enriched in the putative LTR enhancers than to those enriched in the putative LTR promoters of paired ACRs (**fig. S4C, tables S11-14**). Consistently, the single ACRs showed small TF footprints (<40 bp) that were well-aligned with the enriched TF motifs (**Fig. 2F, G, tables S11-14**). Per-fiber chromatin accessibility at these single ACRs was more heterogeneous than that observed at putative enhancers of paired ACRs (**Fig. 2F-I, S4B**). However, the occupancy of small footprints along accessible fibers within these single ACRs was consistent with that of the putative LTR enhancers of paired ACRs (**Fig. 2F-I**).

In stark contrast to paired ACRs in LTRs, or ACRs elsewhere in the maize genome, we observed that a subset of single ACRs in LTRs exhibited hyper-5mCpG methylation coinciding with chromatin accessibility (**Fig. 2B, J, Fig. 3A**), two epigenetic marks thought to be mutually exclusive. Leveraging the single-molecule nature of our chromatin accessibility and 5mCpG methylation calls, we demonstrated that chromatin accessibility and hyper-5mCpG methylation co-occurred and overlapped along the same chromatin fiber at these single ACRs of LTR retrotransposons (**Fig. 2B, table S15**). Hyper-5mCpG methylated ACRs and hypo-5mCpG methylated ACRs showed similar TF footprint occupancy in accessible fibers (**fig. S5B**).

**Fig. 3.**
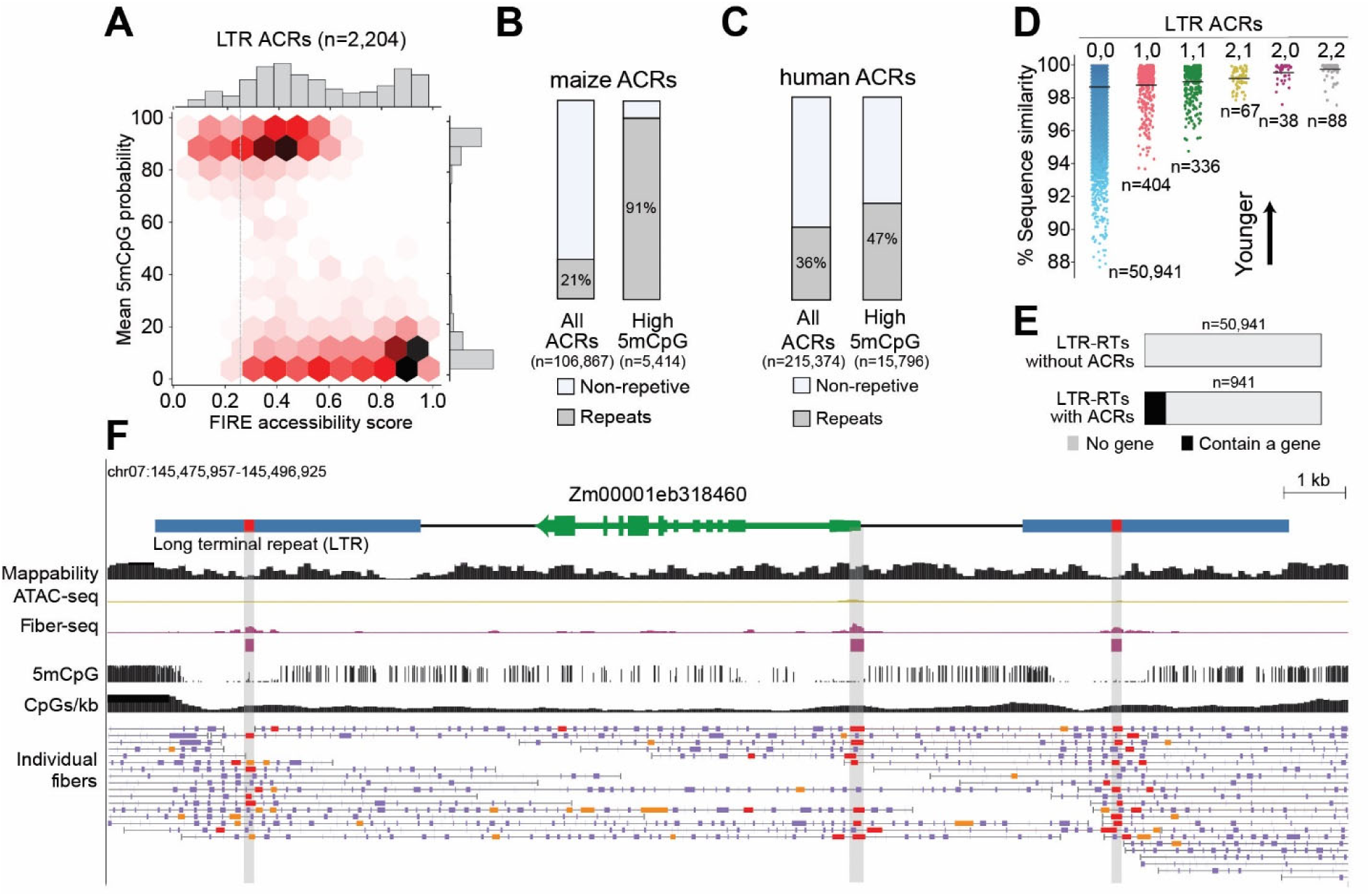
Features of FIRE ACRs in repetitive elements. (**A**) Hexbin plot shows FIRE accessibility scores (X-axis) and mean 5mCpG probabilities (Y-axis) for 2,204 ACRs in LTRs. Shades of red denote frequency, also shown in plotted histograms (top, right). 57% (1261/2204) of FIRE ACRs in LTRs were highly 5mCpG-methylated and 37% (733/1978) of high-confidence FIRE ACRs within LTRs (FIRE accessibility score greater than 0.25, grey dotted line) were highly 5mCpG-methylated. (**B**) Percentage of all FIRE ACRs and FIRE ACRs with high 5mCpG methylation (mean 5mCpG methylation over 50%) that overlap with an annotated repeat by more than 50 bp. (**C**) Fraction of all human FIRE ACRs and human FIRE ACRs with high 5mCpG methylation (mean CpG methylation of over 50%) that overlap an annotated repeat by more than 50 bp. FIRE ACRs calls from human cell line GM12878 (*43*). (**D**) The presence of FIRE ACRs correlates with the sequence similarity of left and right LTRs, a measure of evolutionary age. LTRs with paired bilateral FIRE ACRs showed the greatest sequence similarity while those without FIRE ACRs showed the least. 0,0, no ACRs; 1,0, single unilateral ACR; 1,1, single bilateral ACRs; 2,1, paired ACR in one LTR, single in the other; 2,0, paired ACR in one LTR, none in the other; 2,2, paired bilateral ACRs. Rare instances of other configurations are omitted. (**E**) Fraction of intact LTR retrotransposons with or without at least one FIRE ACR that contain an annotated gene. (**F**) The highly duplicated, well-annotated gene (Zm00001eb318460, green) within an LTR retrotransposon is a candidate for TE-enabled gene amplification. Tracks as in (Fig. 2A). There are single bilateral ACRs present in the LTRs, in addition to a FIRE ACR marking the transcription start site of this gene (highlighted in grey). Statistical analyses and p-values for Fig. 3B**-E** are in **table S1**.

As expected, hyper-5mCpG methylation was rarely seen overlapping FIRE ACRs elsewhere in the maize genome. The rare ACRs with simultaneous hyper-5mCpG methylation and chromatin accessibility were almost exclusively present within repeat elements (**Fig. 3B**), with 15% of these corresponding to single ACRs in intact LTR retrotransposons. In humans, this unexpected co-occurrence of chromatin accessibility and hyper-5mCpG methylation was also rare but not substantially enriched in repeats (**Fig. 3C**). These results point to the plant-specific RNA-mediated DNA methylation pathway as contributing to this unusual co-occurrence of these two epigenetic marks in maize. However, further analysis of the methylation signatures typical of RNA-mediated DNA methylation or other chromatin states at these low-mappability loci was not feasible because the publicly available data sets resulted from short-read sequencing (*25*).

Given the distinct chromatin features of LTRs containing paired and single ACRs, we reasoned that LTR retrotransposons containing the former might be evolutionarily younger TEs, while TEs containing the latter might be older but still younger than the many fully silent transposons. To address evolutionary age, we examined the sequence similarity between the left and the right LTR of each intact LTR retrotransposon as a metric reflecting time since transposition. LTR retrotransposons with exactly one FIRE ACR per LTR (single ACR) showed greater mean sequence similarity than those without FIRE ACRs (99.0% vs 98.7%, p-value=0.004, Mann-Whitney U test) (**Fig. 3D**). LTR retrotransposons with exactly two FIRE ACRs per LTR (paired ACRs) showed a mean sequence similarity of 99.8%, significantly greater than those with one FIRE ACR per LTR (p-value=6.8e-26, Mann-Whitney U test) (**Fig. 3D**). Thus, recently transposed LTR retrotransposons have a characteristic chromatin accessibility and 5mCpG methylation pattern that degenerates with evolutionary age.

## ACRs in LTRs are co-opted as gene promoters and facilitate amplification of host genes

TEs have been long thought to shape host gene regulation by adding or disrupting promoters, enhancers, insulators and coding regions (*25*, *26*). In humans, TE-derived promoters have been inferred via mapping of transcription start sites (*27*) and TF binding sites overlapping TE sequences (*28*, *29*). However, in maize, attempts to infer the regulatory effects of LTR retrotransposons have largely been limited to studying gene expression patterns associated with the presence or absence of neighboring polymorphic TEs that overlap ATAC ACRs (*28*) – an analysis that is severely limited by short-read mappability issues inherent to TEs. We sought to leverage our comprehensive maps of FIRE ACRs across intact maize LTR retrotransposons to identify LTRs that may be shaping host gene regulation (**fig. S6A, B**). Using this approach, we discovered that the putative target gene impacted by LTR FIRE ACRs often resided within the LTR retrotransposon itself. In fact, of the 941 LTR retrotransposons with ACRs in B73, 114 (12%) contained an annotated gene within the intact retrotransposon (**fig. S6C, D**), 24-fold greater than LTR retrotransposons without ACRs (**Fig. 3E**). Of these 114, the promoters of 49 annotated genes were marked by a FIRE ACR, with 48 co-opting one of the LTR ACRs as their promoter (**fig. S6D**). Overall, these findings indicate that a major way that LTRs shape maize host gene regulation is by providing a novel LTR-encoded promoter element.

In one case, the internal gene (Zm00001eb318460) maintained its promoter, marked by a FIRE ACR, in addition to single FIRE ACRs in the flanking bilateral LTRs (**Fig. 3F**). This histone deacetylase complex gene is highly expressed (79% percentile in dark-grown maize leaves) (*30*) and has orthologs in the close maize relative *Sorghum bicolor*, its ancestor teosinte (*Zea mays ssp. mexicana*) (*31*) and other grasses. In human, Alu TEs have been proposed to enable segmental duplication (*32*), so we sought to evaluate whether the gene’s residence within the LTR retrotransposon might be associated with its duplication within the B73 genome. Consistent with this hypothesis, we found numerous Zm00001eb318460 paralogs in the B73 genome (21 amino acid blast hits with e-value<1-e50). This is a highly unusual level of gene duplication for maize genes with a similar expression level and length, as 93% of similar genes showed <5 amino acid blast hits (**table S16**). Taken together, these findings provide evidence that TEs enable gene amplification in maize.

## Diffuse chromatin accessibility marks putative insertion sites of hAT TEs

In general, transposons have been shown to preferentially insert into accessible chromatin both *in vitro* and *in vivo* (*33*, *34*). However, the epigenetic features that predispose genomic loci to insertion of class II (DNA) TEs, in particular hAT TEs, remain unresolved. The first gene reported by McClintock to be susceptible to insertion of hAT TEs is the C locus (*33–37*), now called C1 or colored aleurone 1 gene. The C1 gene body showed unusual diffuse Fiber-seq chromatin accessibility (**Fig. 4A**), ranking among the top 5% of all genes (**Fig. 4B**). We also observed hypo- 5mCpG methylation across the C1 gene body, ranking among the bottom 5% of all genes (**Fig. 4C**). Other genes identified by McClintock as having hAT TE insertions also showed diffuse Fiber-seq chromatin accessibility and hypo-5mCpG methylation within their respective gene bodies (**fig. S7**) (*35–39*). While diffuse gene body chromatin accessibility was not correlated with gene expression (**Fig. 4B**), this feature was strongly correlated with hypo-5mCpG methylation (**Fig. 4C**) (*40–44*). These findings suggest that both diffuse gene body chromatin accessibility and hypo- 5mCpG methylation may uniquely mark preferred hAT TE insertion sites. To test this hypothesis, we identified over 32,000 loci in the B73 maize reference genome that contain hAT TE insertions in exactly one of the 25 non-B73 NAM founder lines (*45*). Indeed, the hAT TE insertion sites were substantially more accessible in B73 than control regions (**Fig. 4D, table S17**) and were preferentially marked by hypo-5mCpG methylation (**table S17**). We thus show that the pervasive epigenetic marks of diffuse Fiber-seq chromatin accessibility and hypo-5mCpG methylation guide the insertional landscape of hAT TEs in maize, including those initially described by McClintock.

**Fig. 4.**
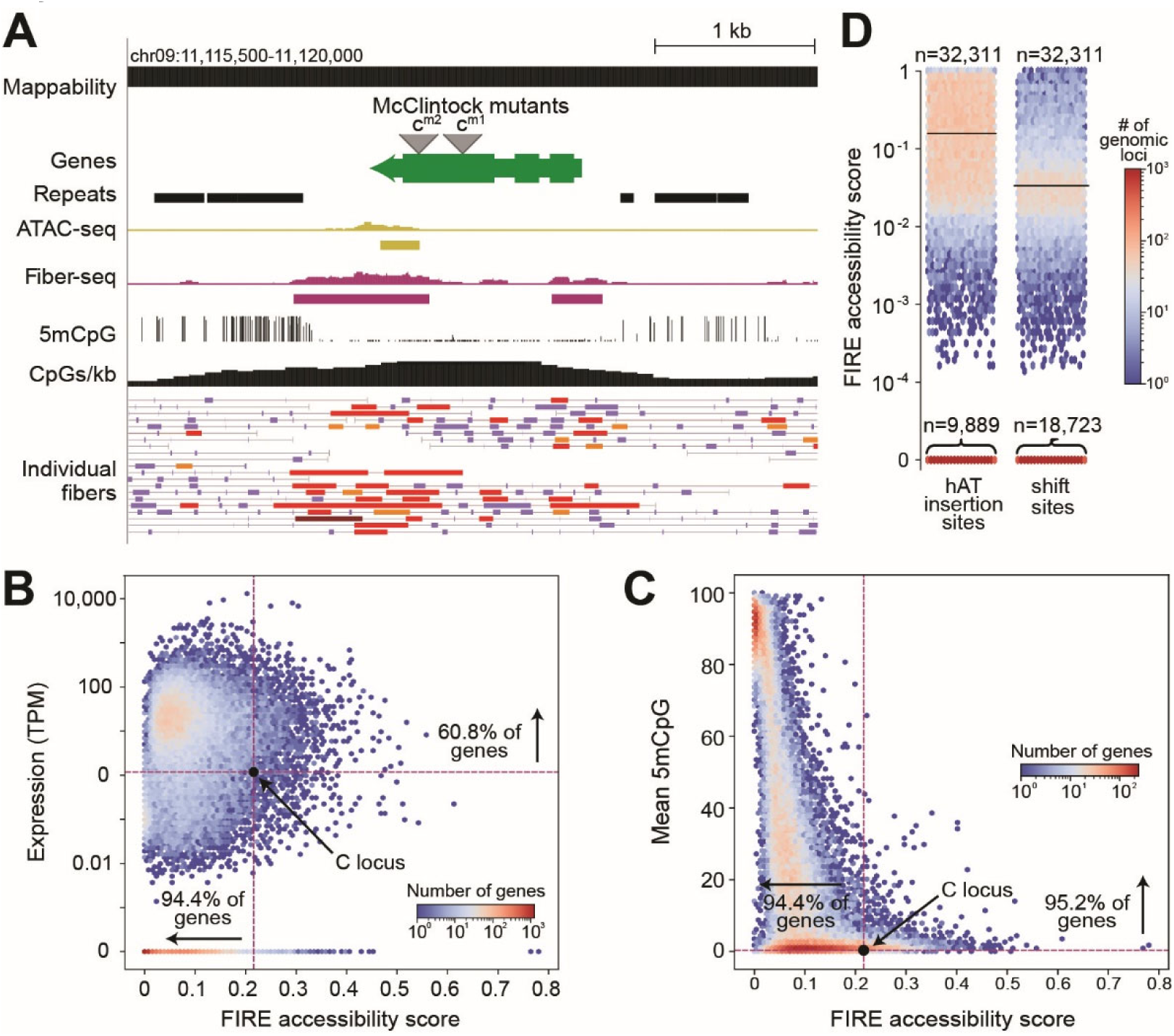
hAT TEs tend to insert in regions with diffuse chromatin accessibility detected by Fiber-seq. **(A)** Screenshot of the locus containing the C1 or colored aleurone 1 gene (green, Zm00001eb373660; called the C locus by McClintock). Tracks as in Fig. 2. hAT TE insertions identified by McClintock as mutant alleles *c^m-1^* and *c^m-2^*. The C1 gene in B73, which does not contain a hAT TE, showed diffuse gene body accessibility detected in Fiber-seq (purple) coinciding with unusual gene body hypo-5mCpG methylation. **(B)** The C1 gene body showed higher FIRE accessibility scores than 94.4% of other genes while showing only slightly above average gene expression. (**C**) The C1 gene body showed lower 5mCpG gene body methylation than 95.2% of genes. (**D**) Loci identified as hAT TE insertions sites in exactly one of the 25 non-B73 NAM strains (*45*) were more likely to show diffuse gene body accessibility (shown as FIRE accessibility scores) than control loci (same-size regions 10 kb shifted to the right in the B73 genome). Mean and median FIRE accessibility scores for hAT TE insertion sites were 0.159 and 0.046. Mean and median FIRE accessibility scores for shifted control regions were 0.035 and 0. Horizontal lines indicate mean FIRE accessibility scores. Thousands of hAT TE insertion sites showed FIRE accessibility scores of 0 (n= 9,889). Statistical analysis and p-value for Fig. 4D are in **table S1**.

## Discussion

The contemporary maize genome is largely comprised of repetitive DNA, with the crop’s ∼40,000 genes clustered in small islands of non-repetitive sequence (*46*). The promise of long-read sequencing for probing regulatory activity in repetitive plant genomes has been illustrated by prior studies in maize (*46*) and Arabidopsis (*47*). However, the reduced accuracy associated with these prior studies’ reliance on single-pass long read sequencing (*48*) has limited their ability to perform an in-depth *de novo* characterization of individual regulatory elements and their associated single-molecule protein occupancy patterns within transposons. Using Fiber-seq paired with PacBio HiFi sequencing in dark-grown maize leaves, we perform *de novo* identification of ACRs in maize, identifying twice as many ACRs as compared to using ATAC-seq in paired samples. This dramatic increase in ACR calls is primarily due to the experimental issues with ATAC-seq for detecting short regulatory elements. The ACRs specific to Fiber-seq share the genomic and epigenetic features of ACRs identified by both methods. Furthermore, they often mark loci that show a strong ATAC-seq signal in other tissues. We attribute this phenomenon of ‘ACR foreshadowing’ to the less frequent and more stochastic TF occupancy at these sites in leaf tissue, which can be captured by Fiber-seq due to its single-molecule resolution. By contrast, the more frequent and more consistent TF occupancy of these same sites in a different tissue results in ACRs that are detectable by both methods.

Unencumbered by mappability concerns, we discover that only 2% (941/51,882) of intact LTR retrotransposons in the maize genome show evidence of regulatory activity, highlighting the efficiency of the plant-specific RNA-mediated DNA methylation pathway to silence the vast majority of transposons in maize. Furthermore, only 94 intact LTR retrotransposons in the maize genome show chromatin accessibility in LTRs in their putative enhancer and promoter elements, chromatin features presumably essential for enabling transposon mobilization.

The presence and specific configurations of ACRs in LTR retrotransposons is correlated with the evolutionary age of that LTR retrotransposon, with accessibility increasingly lost in older transposons. LTR retrotransposons that lose chromatin accessibility preferentially maintain an accessible putative enhancer element. However, the epigenetic pattern at these putative LTR enhancer elements markedly diverges from that of canonical regulatory elements in the rest of the genome. Specifically, on the same chromatin fiber, these enhancer elements can show the unexpected co-occurrence of chromatin accessibility and hyper-5mCpG methylation, two epigenetic marks widely thought to be mutually exclusive. This plant-specific epigenetic pattern raises questions about the nature of the DNA-binding proteins generating accessibility at these sites, which we show can directly occupy these hyper-5mCpG methylated accessible elements. Taken together, these results suggest that plant-specific DNA methylation pathways first suppress the proliferation of LTR retrotransposons by silencing their putative promoters, and subsequently prevent the putative LTR enhancers from being exapted as gene enhancers by hyper-5mCpG methylation. In contrast, we present evidence that a primary mechanism by which LTR retrotransposons may shape maize gene regulation is by creating novel gene promoters from ACRs within the LTRs. We also show an example of an ACR-containing LTR retrotransposon facilitating host gene amplification.

Finally, we find that the loci in which Barbara McClintock discovered TE insertions show unusually low gene body methylation and unusually high gene body accessibility, consistent with the common assumption that gene body methylation protects against TE insertion. While the mechanistic underpinnings of these correlated epigenetic features are unclear, this finding adds to our understanding of the complexity of epigenetics and genome evolution in plants.

## Acknowledgments

We thank Stanley Fields, Olivia Waltner, Michelle Stitzer, Edward Buckler, and Jeffrey Ross-Ibarra for helpful data analysis suggestions and discussion of results as well as detailed manuscript comments.

## Authors contributions

M.O.H., N. A. M., B.R-C., Y.M., E.J.B. and J.R. performed experiments, M.O.H., K.L.B., J.K.M., M.R.V. and T.W. T. performed computational analyses. M.O.H., K.L.B., T.W.T., J.K.M., J.T.C., C.T., C.Q., A.B.S. prepared figures and wrote the manuscript. M.O.H., J.T.C., C.Q, C.T., A.B.S. conceived and designed the experiments.

## Data and materials availability

Raw and processed sequencing data is available from the NCBI Short Read Archive (SRA) under Bioproject PRJNA1119563.

## List of Supplementary Materials

Materials and Methods Figs. S1 to S7

Tables S1 to S17

## Materials and Methods

### Maize mesophyll protoplast generation

We used the PEG transformation method of maize mesophyll protoplasts as described in (*53*). Maize (*Zea mays* L. cultivar B73) seeds were soaked in water overnight at 25°C. The seeds were germinated in soil for 3 days under long day conditions (16 hours light, 8 hours dark) at 25°C, then moved to complete darkness at 25°C for 10-11 days. From each seedling, 10 cm sections from the second and third leaf were cut into thin 0.5 mm strips perpendicular to veins and immediately submerged in 10 ml of protoplasting enzyme solution (0.6 M mannitol, 10 mM MES ph 5.7, 15 mg/ml cellulase R10, 3 mg/ml macerozyme, 1 mM CaCl2, 0.1% [w/v] BSA, and 5 mM beta-mercaptoethanol). The mixture was covered in foil to keep out light, vacuum infiltrated for 3 min at room temperature (RT), and incubated on a shaker at 40 rpm for 2.5 hours at RT. Protoplasts were released by incubating an extra 10 min at 80 rpm. To quench the reaction, 10 mL ice-cold MMG (0.6 M Mannitol, 4 mM MES ph 5.7, 15 mM MgCl2) was added to the enzyme solution and the whole solution was filtered through a 40 µM cell strainer. To pellet protoplasts, the filtrate was split into equal volumes of no more than 10 mL in chilled round-bottom glass centrifuge vials and centrifuged at 100 x g for 4 min at RT. Pellets were resuspended in 1 mL cold MMG each and combined into a single round-bottom vial. To wash, MMG was added to make a total volume of 5 mL and the solution was centrifuged at 100 x g for 3 min at RT. This wash step was repeated two more times. The final pellet was resuspended in 1-2 mL of MMG. A sample of the resuspended protoplasts was diluted 1:20 in MMG and used to count the number of viable cells using Fluorescein Diacetate as a dye.

### ATAC-seq data collection

An aliquot of 50,000 isolated protoplasts was added to new tubes and spun down at 4°C 2000g for 10 min. Supernatant was discarded and the pellet of protoplasts was washed with 750µl of lysis buffer (0.4M Sucrose, 10mM MgCl2, 25mM Tris-HCL pH 8.0, 0.1x Protease inhibitor, 0.5% TritonX). Samples were then spun down at 4°C 1500g for 5min and the supernatant discarded. Samples were then washed once more with buffer (0.4M Sucrose, 10mM MgCl2, 25mM Tris-HCL pH 8.0, 0.1x Protease inhibitor) at 4°C 1500g for 3 min to remove the lysis buffer. The nuclear pellet was then resuspended in 22.5 ddH20 followed by adding 25µl of 2x TD buffer (20mM Tris-HCl pH7.6, 10mM MgCl2, 20% vol/vol DMF) and 2.5µl of Tn5. Samples were then incubated at 37°C for 5 minutes. Reaction was stopped by adding 250µl of Zymo Research DNA Binding Buffer and DNA was purified using Zymo research Clean and concentrator kit. Samples were size selected using 1.8X ampure beads and barcoded with Illumina Nextera Index primers. Final library concentrations were determined using Qubit DNA HS assay and average fragment length was determined using TapeStation D1000 ScreenTape Assay.

### Fiber-seq data collection

1-5 million Isolated protoplasts were spun down at 2000g and resuspended in a 100uL working buffer (400mM sucrose, 15mM Tris-Cl, 15mM NaCl, 60 mM KCl, 1mM EDTA, 0.5mM EGTA, 0.5mM Spermidine), with 1.5uL of 32mM SAM added to a final concentration of 0.8 mM along with 0.5 uL of Hia5 MTase (100U), then carefully mixed by pipetting the 10 times with wide bore tips. Reactions were incubated for 10 minutes at 25℃ then stopped with 3 ul of 20% SDS (1% final concentration) and transferred to a new 1.7 mL microfuge tubes. High molecular weight DNA was then extracted using the Promega Wizard HMW DNA extraction kit A2920. PacBio SMRTbell libraries were then constructed using the manufacturer’s SMRTbell prep kit 3.0 procedure. Two replicate samples were processed.

### Quantification of 6mA/dA by UHPLC-MS/MS

Samples for quantification were treated as previously described (*53*). In brief, 50 ng of DNA from each sample was mixed with 0.02 U phosphodiesterase I (Worthington, cat# LS003926), 1 U Benzonase (Millipore Sigma, cat# E1014), and 2.5 U Quick CIP (NEB, cat#M0525S) in digestion buffer (10 mM Tris, 1 mM MgCl, pH 8 at RT) for 3 hours at 37°C. Single nucleotides were separated from the enzymes by collecting the flow-through of a Nanosep centrifugal filter (MWCO 3 kDa, Pall, cat#OD003C33). The UHPLC-MS/MS analysis of adenosine and m6A was performed on an ACQUITY Premier UPLC System coupled with XEVO-TQ-XS triple quadrupole mass spectrometer. UPLC was performed on a ZORBAX Eclipse Plus C18 column (2.1 × 50 mm I.D., 1.8 μm particle size) (Agilent, cat# 959757–902), 6mA and dA were eluted and separated using 2–50% linear gradient of solvent B (0.1% acetic acid in 100% methanol) in solvent A (0.1% acetic acid in water) within 10 minutes, at a flow rate of 0.3 ml/min. MS/MS analysis was operated in positive ionization mode with 3000 V capillary voltage as well as 150°C and 1000 L/Hour nitrogen drying gas. A multiple reaction monitoring (MRM) mode was adopted with the following m/z transition: 252.10 ->136.09 for dA (collision energy, 14 eV), and 266.2->150.2 for 6mA (collision energy, 15 eV). 6mA and dA were mixed to create standards from 0 to 100nM 6mA, a new standard was measured and used for each run. MassLynX was used to quantify the data.

#### Fiber-seq data processing

Fibertools (*54*) was used to call m6A methylation and label regions as MSPs and nucleosomes on individual reads. Fiber-seq FIRE (*50*) was used to assign FDR values to MSPs and call Fiber-seq ACRs using a FIRE model customized for the maize genome (*55*). Since we trained our model and called FIRE ACRs and elements, the model format used by *fiberseq-FIRE* has changed. To accommodate these changes, a maize model should be re-trained in the future, which can be readily performed and documented in the *fiberseq-FIRE* github repository (*17*). For ACR calling we used the set of peaks identified by the FIRE pipeline with an FDR threshold of 1%. Data for two replicates were combined.

#### ATAC-seq data processing

ATAC-seq read pairs were aligned to the MaizeV5 reference genome (*56*) using bwa v0.7.17-r118 (*57*). The resulting bams were filtered using samtools view (*57*) to discard reads that were unmapped (-F 4), had map quality of zero (-q 1). ACRs were called using MACS2 v2.2.7.1 (*58*), and the narrowPeaks output was merged to generate a non-overlapping set of ACRs. The *ATAC-seq signal* track is a sliding window histogram displaying the number of ATAC read ends, with the height of each 20-bp bar representing the number of Tn5 insertions within a 100-bp window centered on that 20-bp.

#### RNA-seq data used to define expression quantiles and transcription start sites (TSS)

66,143,401 publicly available RNA-seq reads were obtained from NCBI SRA: ERR3322830. These reads were derived from the second leaves of 9-day old, etiolated seedlings (Stelpflug SC et al 2016). Reads were aligned to the maize V5 annotation using hisat2 and counts were tallied using htseq-count. Transcripts Per Million (TPM) were calculated for each gene. 13,542 genes had a TPM of zero. The remainder were split into deciles by expression level, with each decile containing 2991 or 2992 genes. TSS positions were obtained using CAGE data (*59*)

#### Methylation rate (m6A and m5CpG)

For each genomic locus being aggregated, at each 20 bp bin, the number of possible methylation sites was calculated from the individual fiber sequences. The observed methylation events were tallied and divided by the number of possible sites to get a fraction of sites methylated.

#### MSP score and FIRE accessibility score

MSP score is the fraction of fiber-bases within a given region that are annotated as MSPs. FIRE accessibility score is the fraction of fiber-bases within a given region that are annotated as FIRE element (see **fig. S1B**).

#### Percent actuation

For a given genomic region, the number of unique reads (fibers) with at least one FIRE element overlapping the region, divided by the total number of unique reads overlapping the region.

#### Comparing single-cell ATAC-seq to Fiber-seq

The sparse matrix containing binary (cut or no cut) information for all cells from all tissues and all peaks reported in Marand et al 2021 were downloaded (https://ftp.ncbi.nlm.nih.gov/geo/series/GSE155nnn/GSE155178/suppl/GSE155178%5FACR%5Fx%5Fcell.binary.sparse.txt.gz). We then generated a bed file consisting of only peaks with at least one Tn5 insertion in a leaf-designated cell, and reported, for each peak, the fraction of total leaf cells having one or more Tn5 insertion at that site. We used liftOver (*60*) to convert the genomic positions of the peaks in this file from V4 to V5 coordinates, then filtered peaks to retain those that (1) overlap by MACS2 peaks by 100 bp or more, and (2) overlap our FIRE-peaks by 100 bp or more. 39,132 peaks remain. Percent of cells with one or more Tn5 insertion is plotted against percent of fibers containing a fire element (%actuation) in Figure 1g. For this analysis, 58,712 MACS2 narrowPeaks were called on an alignment file (bam) containing 94,945,002 mapped 50-bp paired-end reads. These peaks were merged (bedops -m), resulting in 50,349 MACS2 peaks used above.

#### Short read mappability analysis

2.1 billion fragments were generated evenly distributed across the B73 reference genome chromosomes 1-10. Fragment lengths were sampled from a log-normal distribution fit to one of our ATAC-seq data sets. For each simulated fragment a paired-end read was generated with 50 base reads on either end of the fragment. The true start and end of the fragment was encoded in the read name. We did not simulate per-base errors in these reads; each read matches exactly the reference sequence from which it was generated. These reads were then mapped back to the genome using BWA. The ‘fraction mapped’ for a given region or window was calculated as the number of correctly mapped reads with mapq score > 0 divided by the total number of simulated reads with the outer end (Tn5 insertion) falling in the region. See **fig. S2B**.

#### Annotation of repetitive regions, including all transposable elements

Annotation file Zm-B73-REFERENCE-NAM-5.0.TE.gff3.gz was downloaded from maizegdb.org (*55*)

#### Annotation of regions of the nuclear genome with homology to organellar genomes

Regions of homology within the nuclear genome to organellar genomes were identified as follows for each of the chloroplast and mitochondrial genomes, separately. Paired-end reads were simulated to achieve 100x coverage (142,724 and 579,124 read-pairs, respectively), then mapped to the MaizeV5 reference genome (*56*) using bwa v0.7.17-r118 (*57*). The resulting bams were filtered using samtools view (*61*) to discard reads that were unmapped (-F 4), had map quality of zero (-q 1), or mapped to the centromere (*61*). Alignment files (bams) were then converted to bed files, and overlapping regions were merged.

#### Classification of ACRs

ACRs were sorted hierarchically as follows: (1) all ACRs with low FIRE accessibility score were included in the ‘low FIRE score’ set (medium gray), (2) ACRs with FE length < 200 bp and high FIRE accessibility score were included in the ‘FE length < 200’ set (blue), (3) ACRs with mappability < 80% and both high FIRE accessibility score and FE length >= 200 bp were included in the ‘Low mappability’ set (dark gray), (4) ACRs with high FIRE accessibility score, high FE length, and high mappability are in the ’Regular ACRs’ set (light gray).

#### FiberHMM

Footprints were called on Fiber-seq reads using FiberHMM v1.3.1, as described in Tullius et al. 2024. A new set of transition and starting probabilities were trained using the Maize data, again as previously described.

#### Identifying nearby enhancers

The precise bounds of accessible putative enhancer regions were found using a Gaussian Mixture Model Hidden Markov model (GMM-HMM) to segment regions based on percent nucleosome footprint occupancy. As the positioned mononucleosome footprint found downstream of putative enhancers provided a consistent and clearly defined reference position, the upstream edge of that nucleosome was used to center the footprint occupancy patterns for metaprofiles of the enhancers.

#### Labeling footprints in Fiber-seq

Nucleosome footprints were defined as footprints greater than 90 bp. Putative transcription factor footprints were identified based on a combination of their sub-40 bp size and their overlap with known motifs. Transcription factor footprints overlapping multiple motifs were assigned multiple possible identities, indicated in the corresponding visualization. Putative polymerase footprints were defined based on position relative to the predicted TSS of the promoter, and a size of between 40-80 bp, as previously observed in Tullius et al. 2024 (*62*).

#### Enrichment of GWAS SNPs within different classes of ACRs

SNPs associated with 41 distinct phenotypes (*55*) were used to assess if newly called FIRE ACRs have a similar enrichment of GWAS SNPs to ATAC called ACRs. GWAS SNPs with RMIP<0.05 were removed as described in the paper. FIRE ACRs were split into two categories based on whether they overlap ATAC-seq ACRs as in Figure 2C. For both categories an enrichment was calculated by comparing the fraction of ACR bases covered by GWAS SNPs to the fraction covered in the shifted control category. FIRE ACRs overlapping and not overlapping ATAC-seq ACRs were found to have enrichment values of 3.37 and 3.16 respectively.

#### Calling differential ACRs (dACRs)

ATAC-seq reads from the following six tissues were downloaded from the NCBI Gene Expression Omnibus (Marand et al 2021): Tassel (GSM4696882), Ear (GSM4696883), GSM4696884 (Root1), Axillary_bud1 (GSM4696886), Crown_root1 (GSM4696888), Leaf2 (GSM4696890). For each sample, 100 million read pairs were downloaded, trimmed to 50 bp. Each of the six downloaded samples as well as reads from our in-house dark leaf protoplast sample were aligned to the MaizeV5 reference genome (*56*) using bwa v0.7.17-r118 (*57*). The resulting bams were filtered using samtools view (*61*) to discard reads that were unmapped (-F 4), had map quality of zero (-q 1), or mapped to the centromere (*62*). Because the number of MACS2 peaks is correlated with the number of mapped reads, for each of the seven samples, the number of aligned reads was subsampled to 16M. Peaks were called using MACS v2.2.7.1 (*63*), and the narrowPeaks output was merged to generate a non-overlapping set of peaks for each of the seven samples. A union set of 80,641 peaks was generated by merging the seven sets of peaks (bedops -m) (*64*). TN5 insertions were tallied in each unionpeak for each of the seven samples and per-bp accessibility was calculated by dividing by the peak length. Because our aim was to find differential ACRs that were inaccessible in dark leaf protoplast, we defined differential ACRs as those that (1) had fewer per-bp TN5 insertions than twice the minimum DLP cutcounts in a union peak overlapping a called DLP peak, and (2) the difference between the accessibility of most-accessible sample and the dark leaf protoplast sample was in the 75th percentile or greater. These 2,826 dACRs are in **table S8**.

#### Identification of solo LTRs

LTR sequences from intact LTR retrotransposons containing at least one FIRE ACR within either LTR were aligned to the maize genome using blastn (*64*). Matches with bitscore greater than 1400 and length greater than 1000 bps were retained and merged (bedops -m). Next, we identified matches that (i) did not overlap another intact LTR retrotransposon, and (ii) contained a FIRE ACR. These are listed in **table S10**.

#### Identification of hAT insertion sites

hAT insertion sites are defined as 200-bp windows centered on the location of a hAT transposon polymorphism in which B73 lacks the hAT transposon and exactly one of the 25 NAM lines contains a hAT transposon (*45*).

## Supplemental Tables

**Table S1.** p-values for statistical analyses

**Table S2.** FIRE ACRs

**TableS3.** ATAC ACRs

**Table S4.** chrPt_mappedtonuc.tsv

142,724 pairs of 50 bp reads were simulated from the 142,724 bp length plastid genome, generating 100x coverage. These reads were aligned to a fasta file consisting of the ten nuclear chromosomes, the mitochondrial genome, and the plastid genome. The resulting bam (alignment) file was converted to a bed file (bedtools bamtobed) and overlapping lines were merged (bedops -m).

**Table S5.** chrMt_mappedtonuc.tsv

579,124 pairs of 50 bp reads were simulated from the 579,124 bp length mitochondrial genome, generating 100x coverage. These reads were aligned to a fasta file consisting of the ten nuclear chromosomes, the mitochondrial genome, and the plastid genome. The resulting bam (alignment) file was converted to a bed file (bedtools bamtobed) and overlapping lines were merged (bedops -m).

**Table S6.** Cell actuation vs FIBER actuation

Percent of cells containing at least one Tn5 within this peak vs percent of fibers containing at least one FIRE element overlapping peak.

**Table S7.** Union ATAC ACRs

**Table S8.** Differential ACRs (dACRs)

ATAC ACRs lacking ATAC signal in etiolated leaf protoplasts.

**Table S9.** All intact LTR RTs, with the number of FIRE ACRs contained within the long terminal repeats and the strand information indicated.

**Table S10.** Solo LTRs containing FIRE ACRs

**Table S11.** Motifs enriched in first-of-two paired ACRs vs second-of-two paired ACRs

**Table S12.** Motifs enriched in second-of-two paired ACRs vs first-of-two paired ACRs

**Table S13.** Motifs enriched in single ACR vs first-of-two paired ACRs (putative enhancer)

**Table S14.** Motifs enriched in single vs second-of-two paired ACRs (putative promoter)

**Table S15.** FIRE accessibility score and mean 5mCpG-methylation percentage for all FIRE ACRs within LTRs.

**Table S16.** CompGenes_to_Zm00001eb318460

**Table S17.** B73 coordinates of hAT insertions in exactly one of the other 25 NAM lines and FIRE accessibility scores and methylation values.

---

**Fig. S1.**
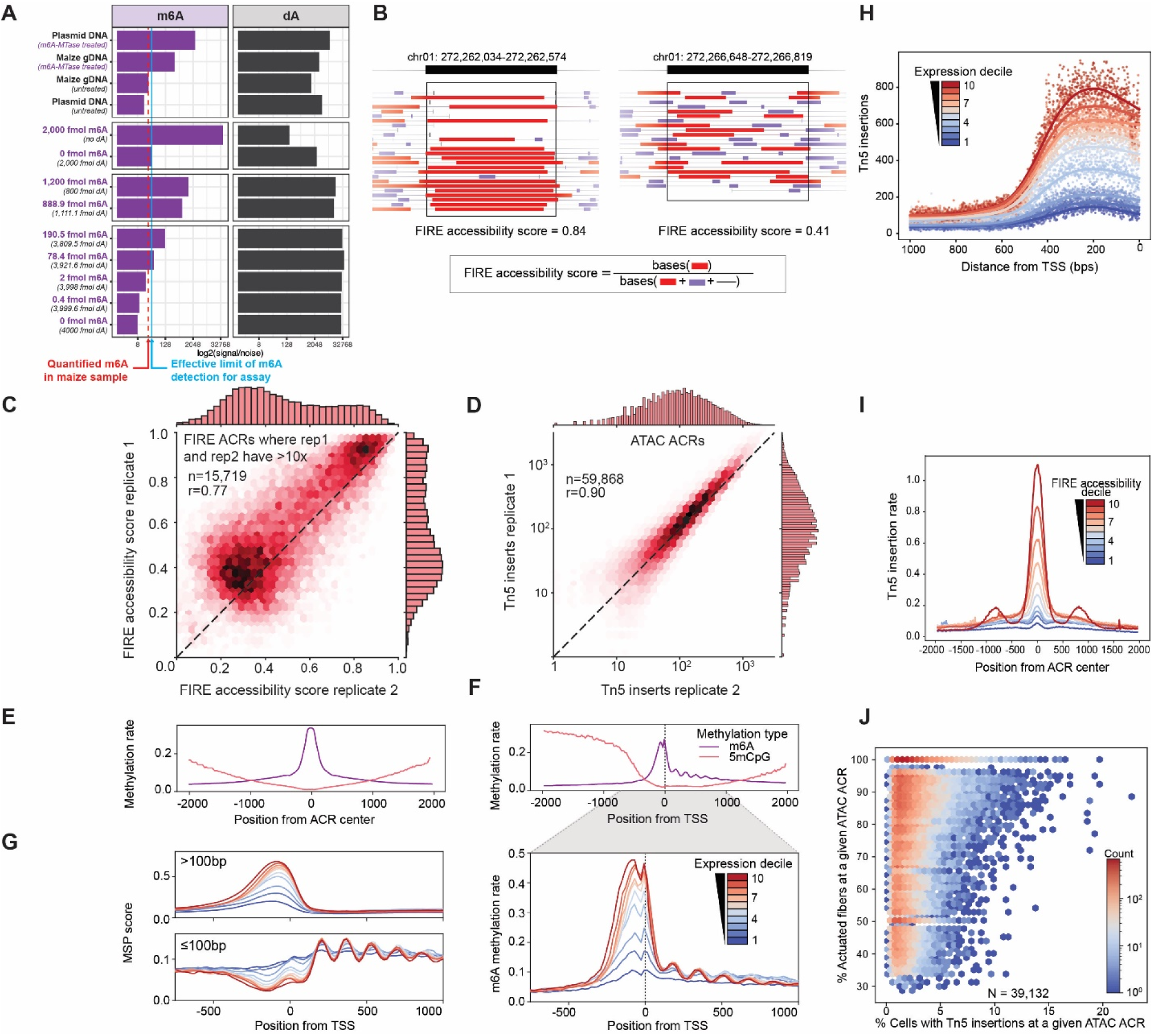
Fiber-seq-derived ACRs show expected patterns at ATAC ACRs and transcription start sites and expected correlation with expression. (**A**) Small molecule mass spectrometry data on nucleotides purified from various samples with known and unknown quantities of m6A and adenine (dA). Shown is the log-scale of the signal-to-noise ratio for each sample. Specifically, this includes plasmid DNA isolated from bacteria that lack any m6A-MTases. As positive controls these samples were treated with a non-specific m6A-MTase prior to nucleotide isolation. For maize, samples were prepared as for Fiber-seq with or without m6A-MTase-treatment. In addition, standards containing defined amounts of m6A and dA were used. The red dashed line corresponds to the signal observed in untreated maize genomic DNA, whereas the solid blue line shows essentially the limit of detection of m6A signal-to-noise for this assay based on the sample that has no m6A injected with a similar amount of total dA. (**B**) Schematic illustrating the calculation of the FIRE accessibility score, a measure of Fiber-seq-derived chromatin accessibility that allows direct comparisons to ATAC-seq-derived chromatin accessibility. Shown are screenshots of two FIRE ACRs with individual fibers showing FIRE elements (red) of different length, in addition to methyltransferase-sensitive patched in purple and non-methylated regions as grey lines. Black boxes mark the respective FIRE ACRs (black bars on top). For any given window, the FIRE accessibility score is calculated as the number of bases annotated as FIRE elements (red) divided by the total number of bases across all fibers mapping within this window (red, purple for methyltransferase-sensitive patches not annotated as FIRE elements, grey for not methylated). FIRE accessibility scores are shown for the two example ACRs. **(C)** Correlation between FIRE accessibility scores for Fiber-seq replicates 1 and 2. Each dot corresponds to ACRs where both replicates have >10x coverage. **(D)** Correlation of Tn5 insertions for union ACRs identified in ATAC-seq replicates 1 and 2. **(E)** m6A methylation peaked at the center of ATAC-seq derived ACRs in paired samples. **(F)** m6A methylation rate peaked immediately upstream of CAGE-defined transcription start sites (TSSs), with phased nucleosomes apparent downstream of TSSs. Average strength of m6A methylation rate upstream of TSSs was monotonically related to expression level of respective downstream genes (expression deciles). The well-phased-nucleosome signal was strongest for highly expressed genes and faded for lowly expressed ones, as expected. (**G**) Methyltransferase-sensitive patches (MSPs) larger than 100 bp constituted the majority of the m6A signal at TSSs, while MSPs shorter than 100 bp showed patterns consistent with well-positioned nucleosomes. MSP scores were calculated in aggregate for each non-overlapping 20 bp window in the region 750 bp upstream and 1 kb downstream of each TSS (see Methods). (**H**) Aggregate plot of Tn5 insertions/base in the 1 kb window upstream of TSSs stratified by downstream gene expression for paired ATAC-seq data, comparable to (**F**). (FIRE elements supporting FIRE ACRs are in red, purple indicates methylation sensitive patches (see Fig.1). (**I**) Aggregate plot of FIRE ACRs stratified into ten deciles based on their FIRE accessibility score. For each FIRE score accessibility decile, the number of Tn5 insertions at each bp within 2 kb of the FIRE ACR center is shown. Accessibility measured by ATAC-seq and Fiber-seq is monotonically correlated. Highly accessible FIRE ACRs tend to show neighboring FIRE ACRs (symmetric signal at highest decile). This signal is in part due to FIRE ACRs in low-mappability LTR retrotransposons (see Fig. 2). (**J**) The single-molecule method Fiber-seq outperforms single-cell ATAC-seq as a quantitative measure of chromatin accessibility. 39,132 ACRs were identified as shared FIRE ACRs in dark-grown maize leaves and ATAC ACRs in a pseudobulked leaf sample (GSM4696890) from Marand et al 2021 (*50*). The percentage of cells containing at least one Tn5 insertion within a shared ACR (% cells accessible) is compared to the percentage of actuated fibers (*i.e*., with a called FIRE element, % actuated Fibers within a given ATAC ACR) underlying the same shared ACR. Each dot represents one shared ACR. Hexbin color reflects the number of dots.

**Fig. S2.**
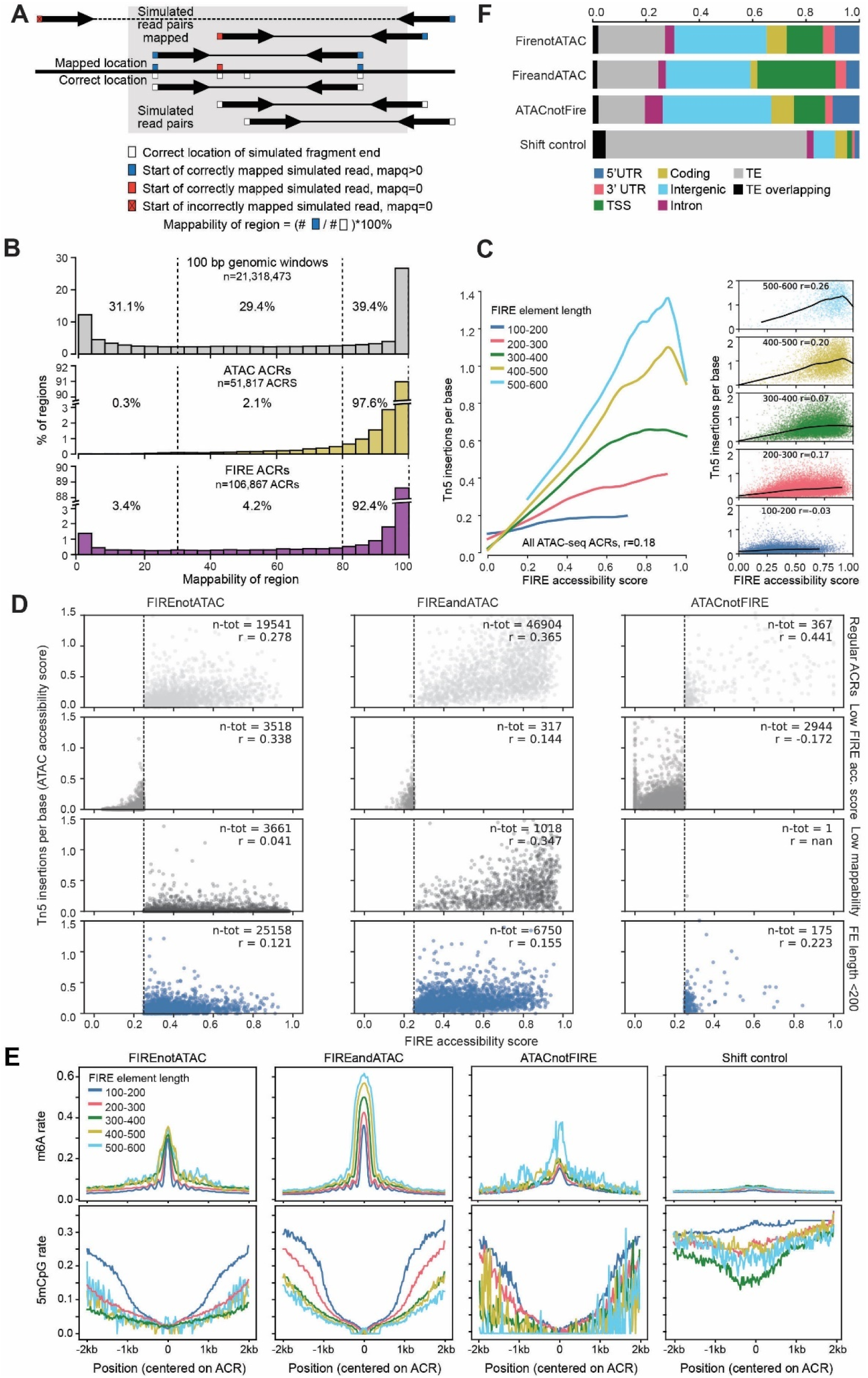
Novel FIRE ACRs comprised of short FIRE elements are bona fide regulatory elements. **(A)** Schematic describing short-read simulation and mappability calculation. We generated 2.1 billion fragments evenly distributed across the B73 reference genome chromosomes 1-10 (see Methods). For each simulated fragment, 50 bp paired-end reads were generated (indicated with thick black arrows). Each read matched exactly the reference sequence from which it was generated. These simulated reads were then mapped back to the genome using BWA. The ‘fraction mapped’ for a given region or window was calculated as the number of correctly mapped reads with mapq score > 0 divided by the total number of simulated reads with the outer end (Tn5 insertion) falling in the region. Mapq scores are indicated by blue and red boxes, incorrectly mapped simulated read shows X in red box (top row). Mappability of regions was determined as percentage of correctly mapped reads with mapq>0. (**B**) Histograms of mappability as in (**A)** for all 21,318,473 non-overlapping 100 bp windows in the maize genome (top panel, grey), 51,817 ATAC ACRs (middle panel, gold), and 106,867 FIRE ACRs (bottom panel, purple). Low mappability explains only in part why Fiber-seq detects many more ACRs than ATAC-seq. (**C**) FIRE ACRs comprised of short FIRE elements are not detected by ATAC-seq. Correlation between FIRE accessibility scores and Tn5 insertions/ base (chromatin accessibility as measured by ATAC-seq) for FIRE ACRs comprised of FIRE elements of indicated length (see inset for legend). **Left**, LOWESS curves fitted to FIRE ACRs in respective length categories. **Right**, plots showing individual values for FIRE ACRs belonging to the five length categories. (**D**) FIRE accessibility score by Tn5 insertions/base (*i.e.*, ATAC accessibility score) for ACRs stratified into 12 categories. Each dot represents an ACR with the labeled row and column properties. As the row categories are overlapping, ACRs were sorted hierarchically as follows: all ACRs with low FIRE accessibility score were included in the ‘low FIRE acc. score’ rows; ACRs with FE length < 200 bp and high FIRE accessibility score were included in the ‘FE length <200’ rows; ACRs with mappability < 80% and both high FIRE accessibility score and FE length >=200 bp were included in the ‘Unmappable’ rows. (**E**) FIRE ACRs that do not overlap with ATAC ACRs show similar patterns of the m6A signal (top) and the 5mCpG signal (bottom) as FIRE ACRs that overlap with ATAC ACRs. Shifted control regions do not display these properties. FIRE element length underlying FIRE ACRs is indicated as in (**C**). (**F**) FIRE ACRs that do not overlap with ATAC ACRs show a similar distribution across genomic compartments as FIRE ACRs that overlap with ATAC ACRs. Statistical analyses and p-values for **fig. S2F** are in **table S1**.

**Fig. S3.**
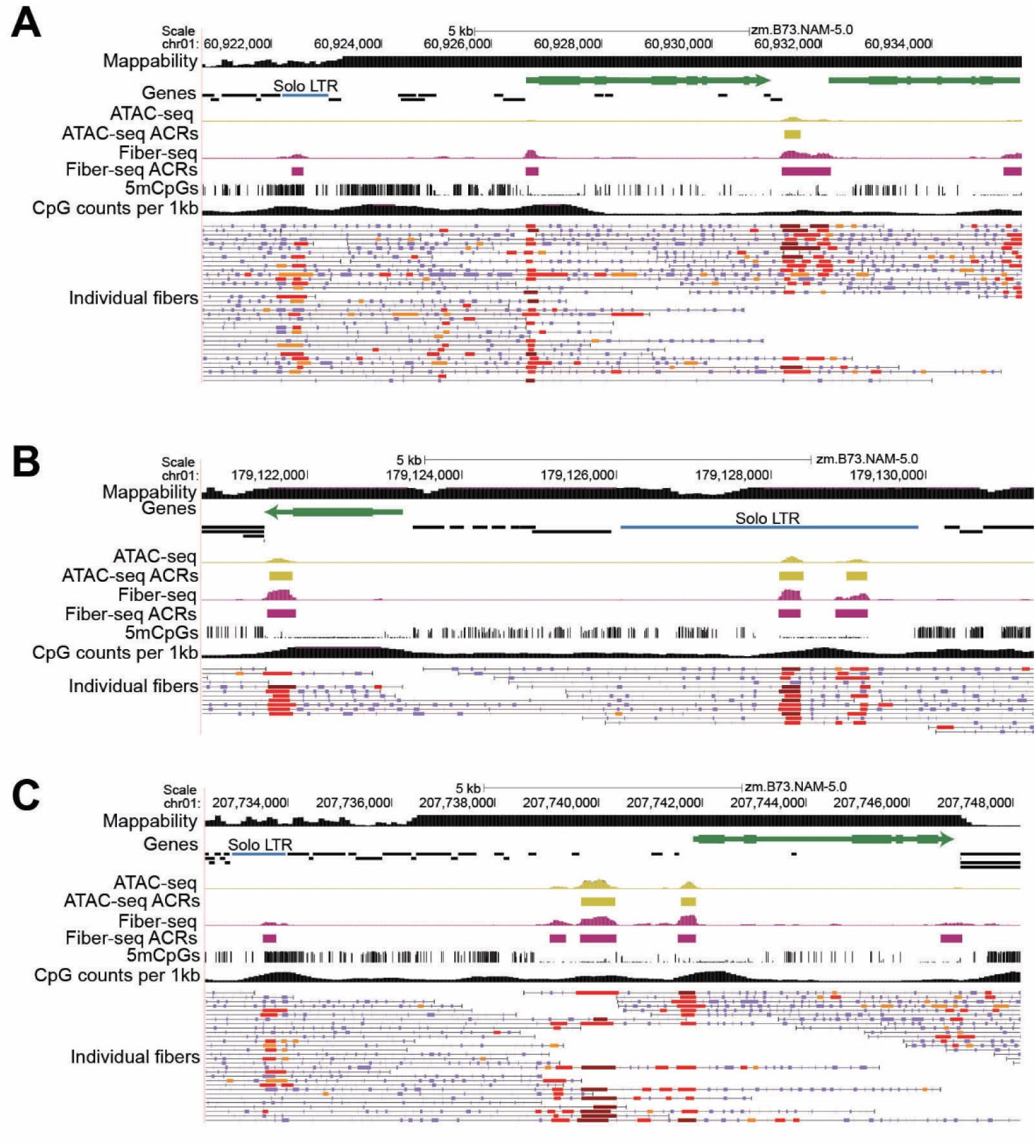
Examples of solo LTRs containing FIRE ACRs. **(A-C)** Solo LTRs containing FIRE ACR are colored blue. **(A)** [chr01:60,920,594-60,935,475] (**B**) [chr01:179,120,635-179,131,399] **(C)** [chr01:207,732,409-207,748,141]. See **table S10** for a comprehensive list.

**Fig. S4.**
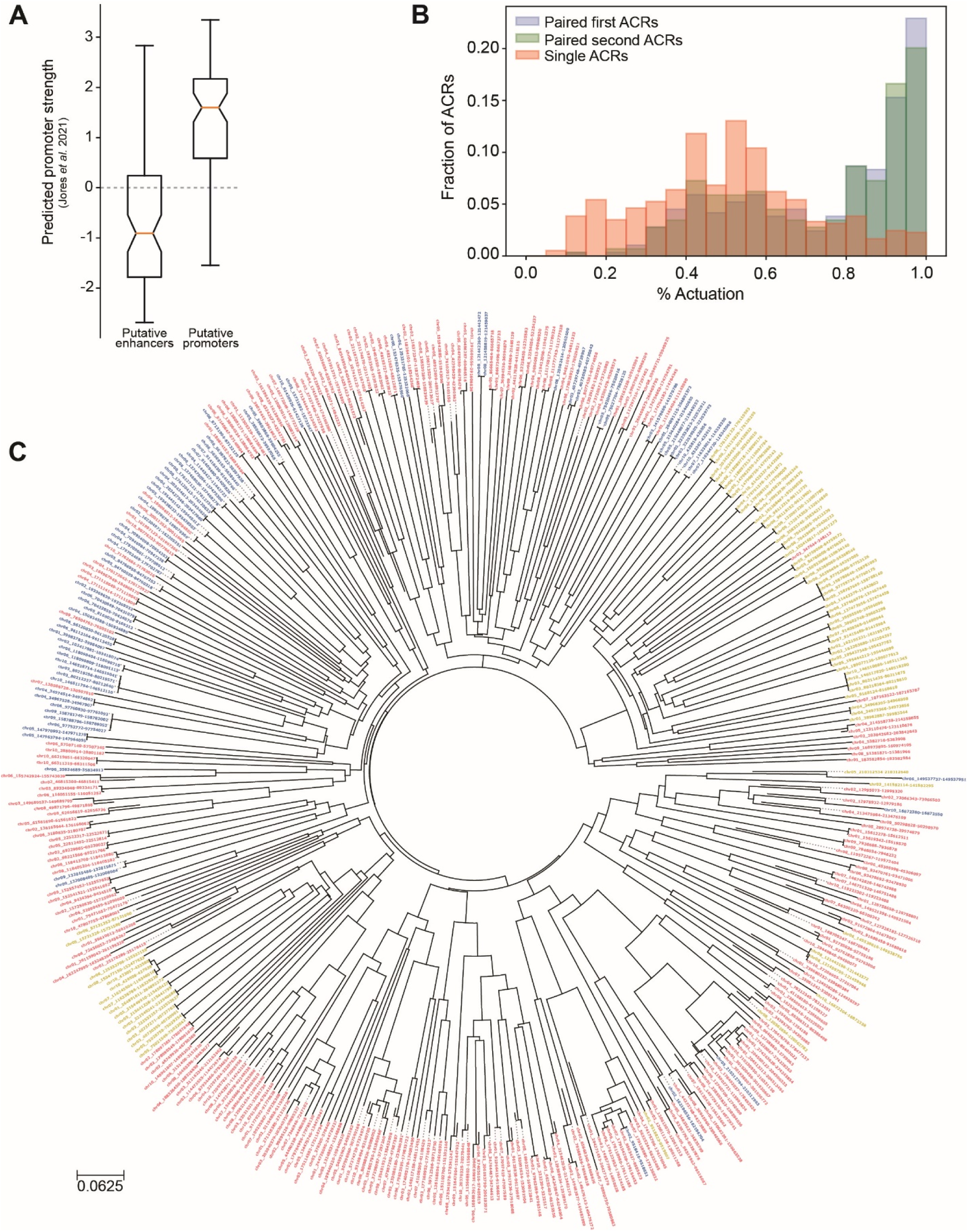
Features of FIRE ACRs within LTR retrotransposons. (**A**) For each putative enhancer-promoter pair, a sequence starting at the 5’ end of the putative enhancer ACR and ending at the 3’end of the putative promoter ACR was extracted. The boxplot shows the predicted promoter strength of the first (coinciding with the putative enhancer) and last 170 bp (coinciding with the putative promoter) of this window. Predictions were made with a CNN model trained on Plant-STARR-seq data for ∼ 75,000 TSS-proximal ACRs (170 bp in length) from Arabidopsis, maize, and sorghum (*51*)**. (B)** Histograms for the percentage actuation (*i.e.*, the percentage of fibers with a FIRE element that comprise a FIRE ACR) for the first of two paired ACRs (putative enhancers), the second of two paired ACRs (putative promoters), and single ACRs. (**C**) Phylogeny of LTR ACRs. Branch length units are in estimated substitutions per site (*52*). Colors indicate ACR types with blue denoting paired first ACRs (putative enhancers), yellow denoting paired second ACRs (putative promoters) and red denoting single ACRs in LTRs.

**Fig. S5.**
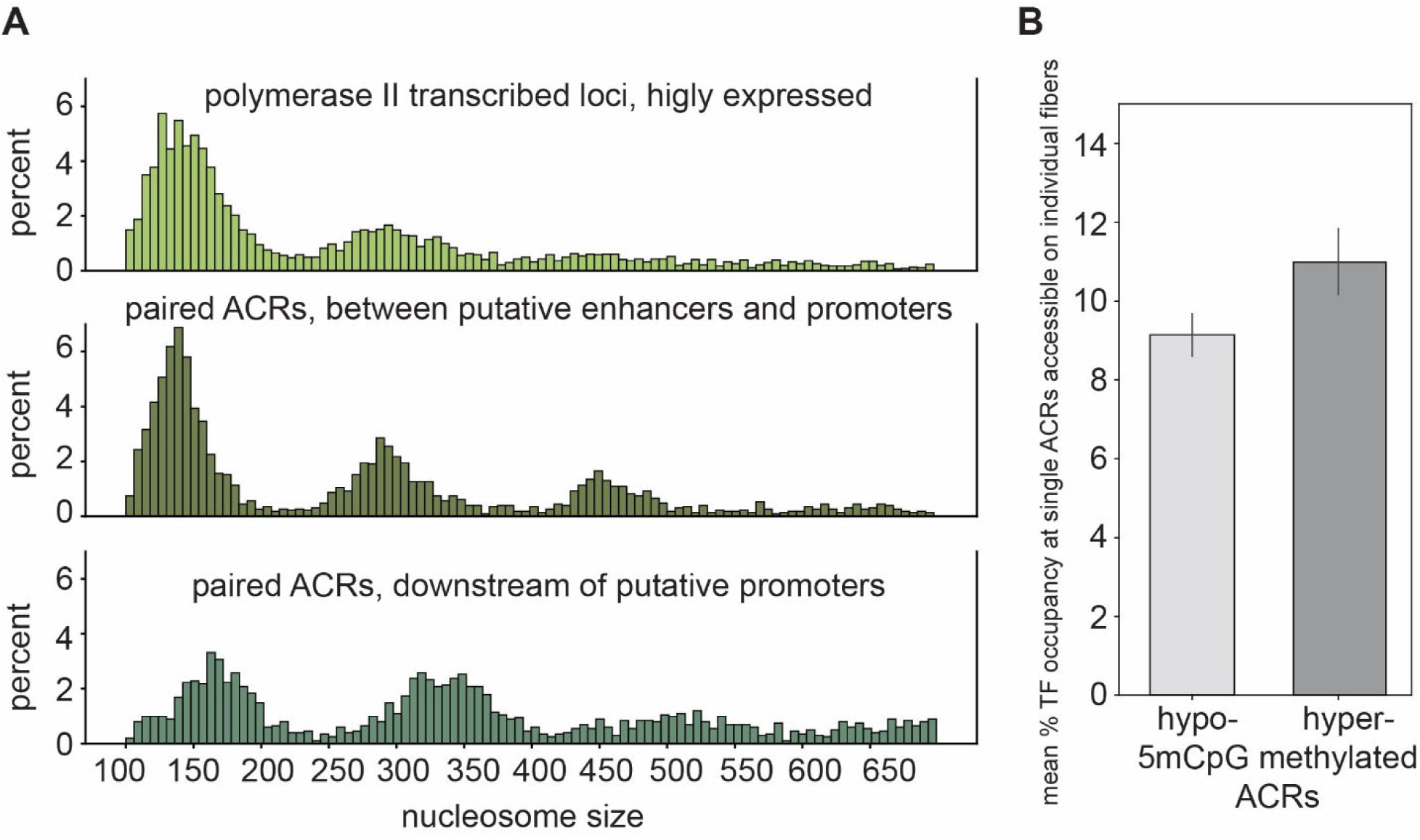
Nucleosome and TF footprint features of LTR FIRE ACRs. (**A**) Histogram showing the percent of footprints sized between 100 and 700 bp identified (top) 100 to 1100bp downstream of the TSS of Pol II genes in the top 3 deciles of expression, (middle) between putative enhancer/promoter pairs, and (bottom) 300-1300 bp downstream of the putative promoter. (**B**) Bar plot showing the mean percent putative TF (10-40 bp) occupancy within +/- 100bp of the center of all hypo- or hyper-methylated single ACRs calculated from reads where the center position of the ACR was not occluded by a nucleosome. Error bars represent the 95th percentile range calculated from 10,000x bootstrapped resampling of each group of reads. Statistical analysis and p-value for **fig. S5B** are in **table S1**.

**Fig S6.**
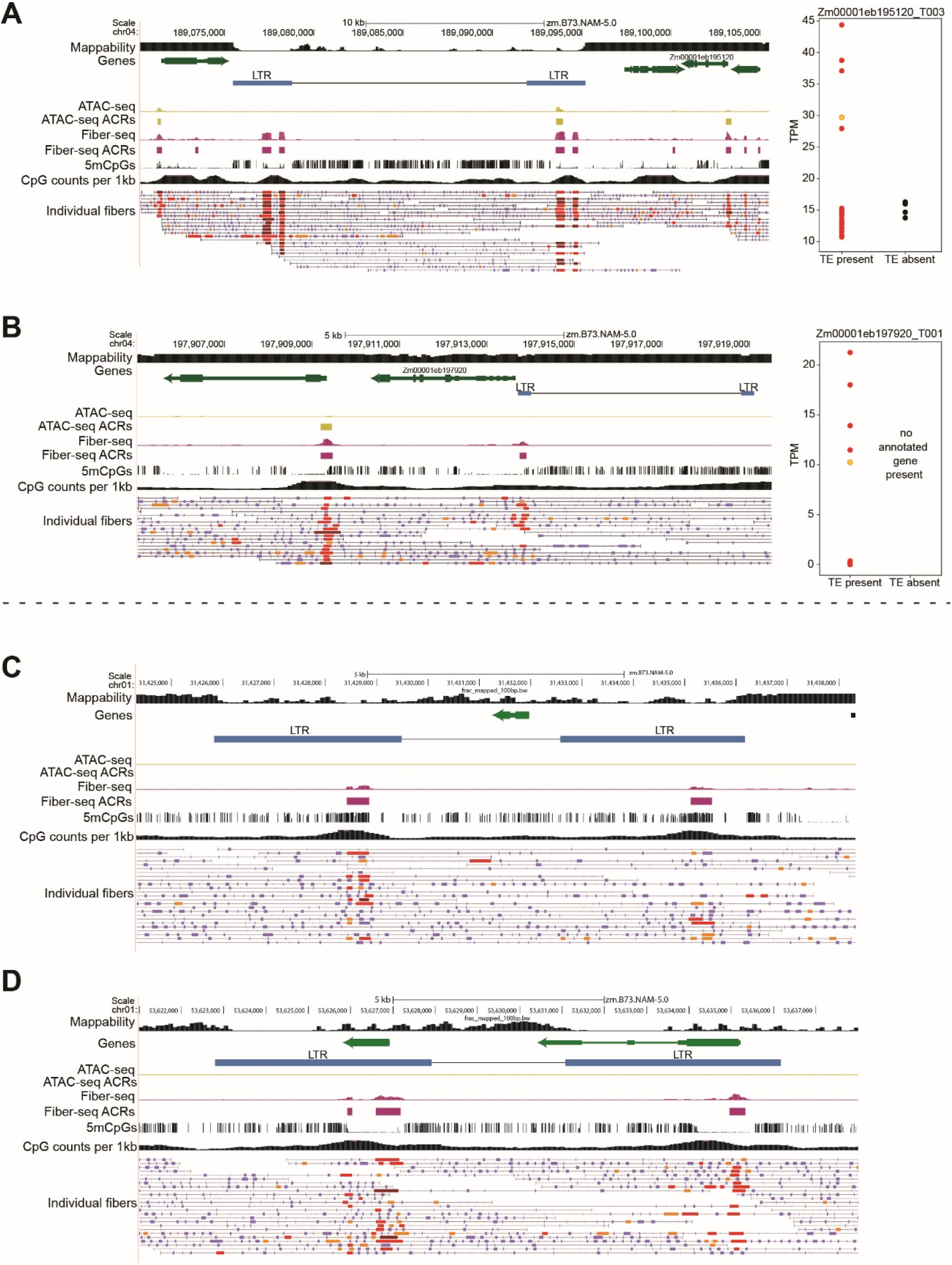
Examples of ACRs in intact polymorphic LTR retrotransposons (A, B) and LTR retrotransposons with non-TE internal genes (C, D). (**A**) Left, intact LTR retrotransposon with blue LTRs is absent in NAM lines: Il14H, Ki3, M37W, P39. Tracks in screenshot as in Fig. 2. Right, expression level of indicated gene in lines with and without the TE, B73 is labeled in yellow. (**B**) Left, intact retrotransposon with blue LTRs is absent in NAM lines: B97, CML228, CML52, Ki11, Ky21, Mo18W, P39. Tracks in screenshot as in Fig. 2. Right, expression level of indicated gene in lines with and without the TE, B73 is labeled in yellow. (**C**) Example of an intact LTR retrotransposon containing one annotated gene between the LTRs and lacking an ACR at the transcription start site. (**D**) Example of an intact LTR retrotransposon containing two annotated genes. For each gene, transcription begins at a FIRE ACR within the LTR.

**Fig. S7.**
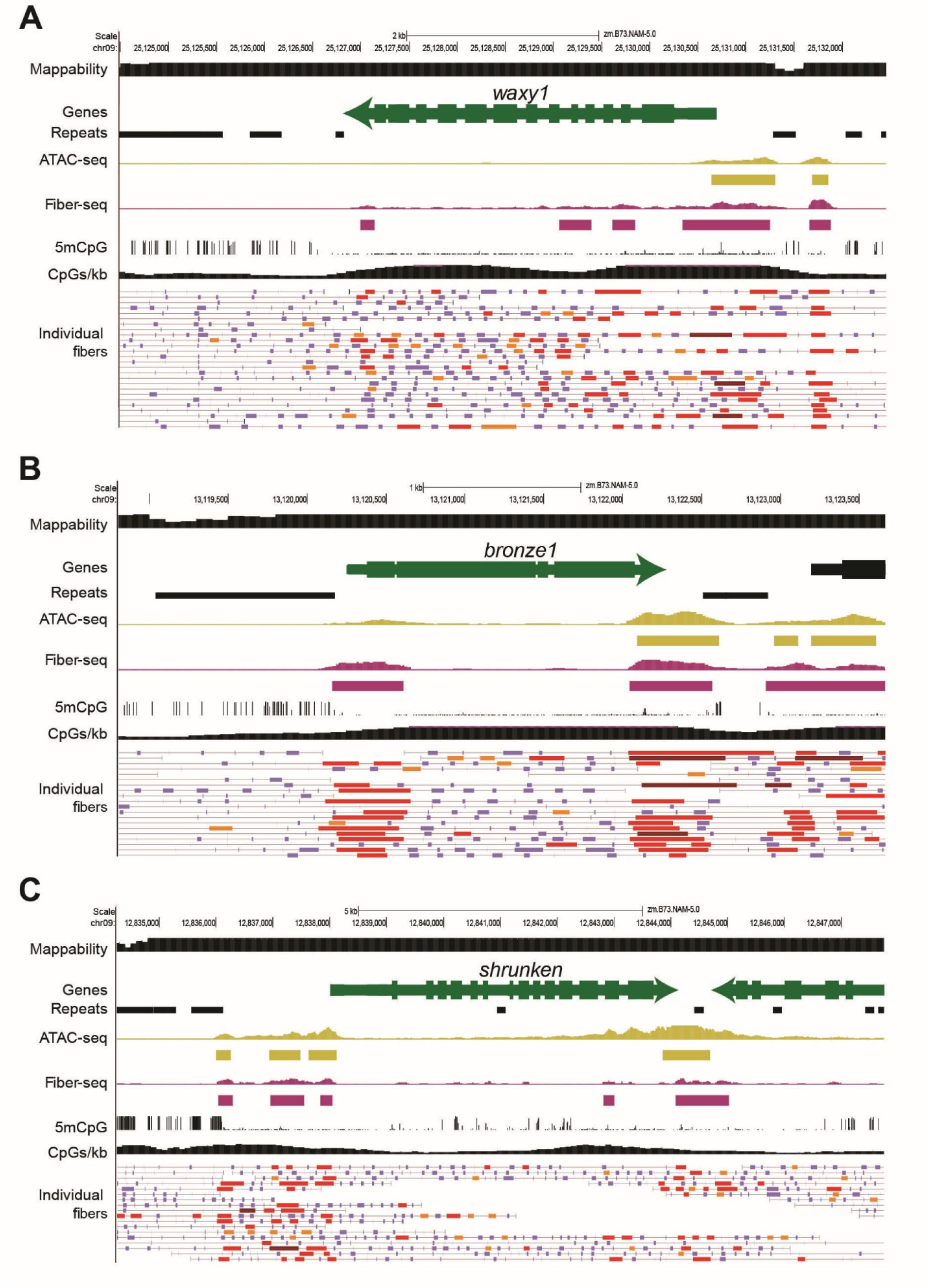
Diffuse chromatin accessibility and hypo-5mCpG methylation is observed at loci discovered as hAT TE insertion sites by McClintock (*51*). **(A)** *waxy1* (Zm00001eb378140; chr09:25,127,146 - 25,129,800), one of the first genes identified by McClintock as having a hAT TE insertion, shows higher gene-body chromatin accessibility than 84.4% of other genes. McClintock identified alleles *Ds wx-m9*, *Ds wx-m6*, *Ac wx-m9*, with the *Ds* or *Ac* prefix indicating whether it was a nonautonomous or autonomous hAT TE, respectively. **(B)** *bronze1* (Zm00001eb374230; chr09:13,118,806-13,123,664), one of the first genes identified by McClintock as having a hAT TE insertion. McClintock identified the *Ac bz-m2* allele. The *Ac* prefix indicates insertion of an autonomous hAT TE. **(C)** *shrunken* (Zm00001eb374090; chr09:12,836,508-12,845,499), one of the first genes identified by McClintock as having a hAT TE insertion. McClintock identified two germinally-stable alleles, *Ds-4864A* and *Ds-5245*, that were “genetically indistinguishable and located just distal to the Shrunken (Sh) locus on the short arm of chromosome 9” and three germinally-unstable alleles, *sh-m6233*, *sh-m5933*, *sh-m6258*, that contain rearrangements at the Sh locus related to a hAT insertion, one of which contains a Ds-mediated 30 kb insertion (*52*). The *Ds* prefix indicates insertion of a nonautonomous hAT TE.

